# A hierarchical multiscale model of forward and backward alpha-band traveling waves in the visual system

**DOI:** 10.1101/2024.11.15.623743

**Authors:** Jakob C. B. Schwenk, Andrea Alamia

**Affiliations:** Centre de Recherche Cerveau et Cognition (CerCo), CNRS, Université de Toulouse, Toulouse, France

## Abstract

Recent studies have shown that low-frequency oscillations in the cortex are often organized as traveling waves. The dynamical properties of these waves, that span different scales, have been linked to both sensory processing and cognitive functions. In EEG recordings, alpha-band (∼10Hz) traveling waves propagate predominantly in both directions of the occipital-frontal axis, with forward waves being most prominent during visual processing, while backward waves dominate at rest and during sensory suppression. While a previous study has proposed a functional model to explain their generation and propagation, a multi-scale, biologically plausible implementation is still lacking. Here, we present a multi-scale network model with mean-field dynamics that, building on known interlaminar and cortico-cortical projections, reproduces the dynamics of alpha-band traveling waves observed in EEG recordings. We show that scalp-level forward and backward waves can arise from two distinct sub-networks that are connected in infragranular layers at each area. At rest, the network generates spontaneous backward waves and switches to a forward state upon bottom-up sensory stimulation, reproducing the dynamics we observed in EEG recordings in healthy participants.. We then expand our model to a cortico-thalamic network with a parallel feedforward pathway through the pulvinar. Our results show that this pathway biases the cortical dynamics to the forward state and that high pulvinar engagement leads to spontaneous forward waves without external input. This result is in line with previous studies suggesting a key role for the pulvinar in directing information flow in the cortex, and provide a computational basis to investigate the role of the pulvinar in cortical dynamics. In summary, our model provides a biologically plausible architecture for modeling the dynamics of macroscale traveling waves. Importantly, our study bridges the gap between distinct scales by connecting laminar mean-field activity to spatial patterns at the scalp level, providing a biologically grounded and comprehensive view of the generation and propagation of alpha-band traveling waves.

## Introduction

Oscillatory activity is a ubiquitous feature of neural processing throughout the brain and has been linked to a broad array of sensory, cognitive, and motor functions (Buzsaki, 2006). The most dominant temporal frequency band in the visual system is the alpha rhythm (approx. 7 – 13 Hz), which has been linked to several cognitive processes. For example, previous studies considered alpha oscillations as a rhythm mainly involved in modulating top-down inhibitions in sensory-specific cortical areas (Jensen & Mazaheri, 2010; Klimesch, 2012; Mathewson et al., 2011; Sadaghiani & Kleinschmidt, 2016). However, due to its strong association with sensory input and phasic influence on perception, alpha oscillations have also been proposed as an internal temporal reference frame for neural processing in the visual system (Busch et al., 2009; Dugué et al., 2011; VanRullen, 2016). More recently, as studies on neural oscillations shift their focus also to consider their spatial dimension, it was demonstrated that alpha oscillations propagate as traveling waves, as measured at the cortical surface (Zhang et al., 2018), as well as using EEG (Lozano-Soldevilla & VanRullen, 2019; for early investigations see also Nunez et al., 2001, and Ito et al., 2005). At the scalp level, the propagation direction of these waves is concentrated mainly on the anterior-to-posterior axis, with waves traveling either forward (FW, i.e., towards anterior sensors) or backward (BW, i.e., in the opposite direction). At rest, as well as during attentional suppression of visual input, the dominant direction is BW, while FW waves are more prominent during visual stimulation and attention (Alamia et al., 2023; Ito et al., 2005; Lozano-Soldevilla & VanRullen, 2019; Nunez et al., 2001; Pang (庞兆阳) et al., 2020). These findings and the previous literature on alpha oscillations suggest that alpha-band traveling waves may play a functional role in the visual system. Yet the cortical spatiotemporal patterns of activity that correspond to FW and BW waves at the scalp level remain unclear. On the one hand, from an experimental point of view, some studies have attempted to infer the cortical sources using M/EEG signals (e.g., Nunez et al., 2001; Zhigalov & Jensen, 2023), but the precision of these methods remains limited and, ultimately, invasive, multiscale recordings (i.e., simultaneous from the scalp and cortex) may be needed to gain a full understanding of the waves’ cortical origins. An increasing number of studies reporting traveling waves (including in the alpha band) from intracortical and cortical surface recordings (Bahramisharif et al., 2013; Bhattacharya et al., 2022; Halgren et al., 2019; Zhang et al., 2018) further highlight the need to bridge the gap between this literature and the EEG findings. From a computational perspective, a relatively simple hierarchical network model could predict the FW/BW waves’ direction in EEG recordings (Alamia & VanRullen, 2019). More specifically, in this model, based on the theoretical framework of predictive coding (Rao & Ballard, 1999), each area in the hierarchy continuously predicts the input it receives from the previous area. Alpha traveling waves then arise naturally from the resulting inter-areal feedback loop (at plausible delays), with FW and BW directions following from input to the first and last stages of the hierarchy, respectively. While this model offers a computational framework for investigating traveling waves at the scalp level, it fails to provide a physiologically plausible architecture that describes the mechanisms involved in generating alpha traveling waves at the level of cortical neural circuits.

In this study, we aimed to bridge this important gap in the literature and provide a biologically plausible model to investigate the cortical generation and propagation of alpha-band traveling waves observable at the scalp level. In implementing our model, we attempted to integrate a hierarchical architecture, as proposed by previous computational studies and theoretical work, into the known connectivity of the visual cortex and its features. First, we designed a model such that the interlaminar and interareal circuits generate forward and backward waves through different pathways and validated the results of our model with human EEG recordings. We then explored how the thalamus could modulate these waves. Specifically, we considered the pulvinar, which is a critical, higher-order thalamic nucleus with projections both from and to the cortex (Shipp, 2003), and also a known generator of alpha-band oscillations (Halgren et al. 2019). Importantly, several studies point to the pulvinar as being involved in the modulation of cortico-cortical communication, including through oscillatory dynamics (Fiebelkorn & Kastner, 2019; Jaramillo et al., 2019; Quax et al., 2017; Saalmann et al., 2012; Zhou et al., 2016). This involvement suggests the pulvinar may also be crucial in generating alpha-band traveling waves, and we explored its effect on cortical waves in our implementation. All in all, the resulting network allows us to accurately describe both the intrinsic generation of alpha -rhythmicity its propagation along the cortical hierarchy, faithfully replicating the propagation patterns observed in EEG recordings. Our results reveal that large-scale FW and BW traveling waves can arise from specific and distinct cortico-cortical connections, that can be modulated by the pulvinar and that their propagation direction is determined by the presence of sensory input.

## Results

We designed a model reproducing the spatial patterns of alpha traveling waves, as observed in EEG recordings, during rest, and in response to visual stimulation. Our implementation is grounded in a few well-known visual system features. First, the visual cortex is organized hierarchically, with largely separate feedforward and feedback pathways (Felleman & Van Essen, 1991; Markov et al., 2014; Vezoli et al., 2021). Another prominent feature is its laminar organization, reflecting both in the interareal flow of feedforward and feedback signals and in the circuit within each area where these signals are combined (Shipp, 2007; Thomson & Bannister, 2003). In addition, there is also strong evidence for a laminar separation of oscillatory sources, as alpha generators have been specifically localized in deep (infragranular) layers (Maier et al., 2010, 2011; Mendoza-Halliday et al., 2024). We considered all of this evidence to constrain the cortical connectivity in our model. In the following, we will describe the model’s structure, characterize its general response behavior, and compare it to experimental data from human EEG recordings during rest and visual stimulation. We then continue with a more detailed analysis of the network’s pathways that generate FW and BW waves. Finally, we explore how the model’s dynamics change by adding a cortico-thalamo-cortical pathway through the pulvinar, investigating how thalamic engagement could reproduce the effects of attentional modulation.

### Model architecture

Our model represents a generalized description of hierarchical processing streams in the visual system without modeling specific cortical areas. The general architecture is illustrated in Figure 1a. It describes a set of *N_Cx_* hierarchically connected cortical areas with identical intra- and interlaminar connectivity. Each area comprises a set of nodes with mean-field dynamics across three laminar compartments: the supragranular layers 2/3, the input layer 4, and the infragranular layers 5/6. Incoming feedforward signals enter each area at L4 (L4_X_), where initial response adaptation is applied through a local inhibitory population (L4_IN_), and are then passed to the supragranular layers (SG_X_) within the same area. From here, two separate connections extend to 1) an ensemble of excitatory and inhibitory supragranular nodes generating the feedforward signal to the next area (SG_E_ and SG_IN_) and 2) deep infragranular layers (IG_IB_) within the same area. The infragranular node in each area (IG_IB_) is modeled as a population of intrinsically bursting neurons that generate a steady oscillation at ∼9 Hz at baseline, based on neurophysiological evidence demonstrating the presence of strong alpha pacemakers in infragranular layers (Steriade et al., 1990; Sun & Dan, 2009). These nodes are unidirectionally connected through excitatory feedback between areas. For its role in generating BW-directed traveling waves in the network (see below), we label this stream of backward-coupled IG_IB_ nodes the *BW-pathway*.

**Figure 1.**
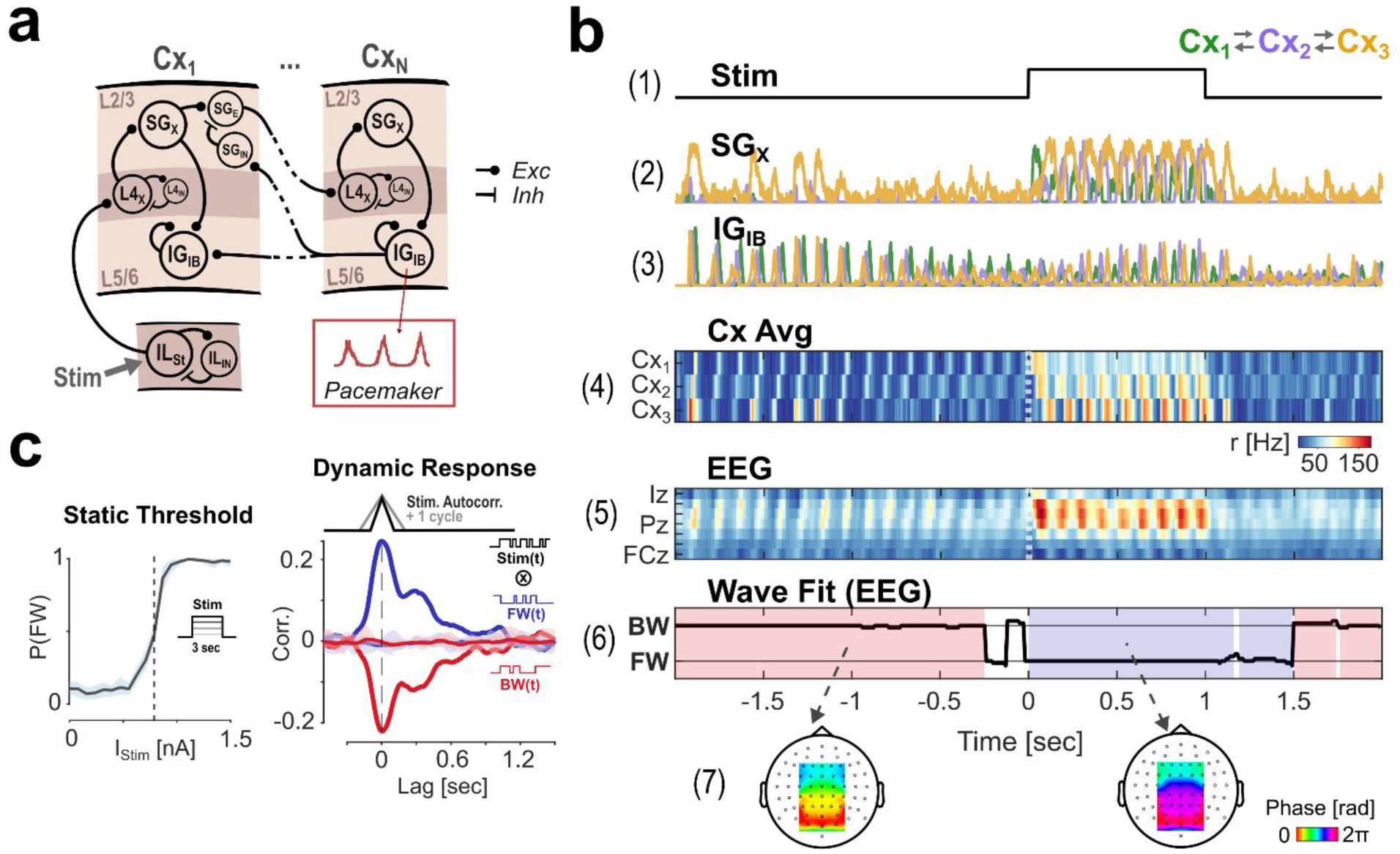
Model overview and basic response behavior. A: Modular architecture. The model comprises a laminar network in a hierarchical stream of cortical areas Cx_1_ to Cx_N_. Each area sends feedforward signals from supragranular layers and receives feedback to both laminar compartments from the infragranular layer of the higher area. Nodes in infragranular layers are pacemakers generating a continuous alpha rhythm. B: Simulated responses and wave fits for a single model run with N_C__x_ = 3 areas during rest and visual stimulation; (1) Stimulus profile (1nA DC step function); (2, 3) cortical activity in supra- and infragranular layers (colors represent different cortical areas); (4) average of all layers’ activity in each cortical region at the source- and (5) at the scalp-level (via forward projection to simulate an EEG signal); (6) traveling wave direction fit: the time-course shows the angle of the best fit to the EEG phase gradient (see Methods for details); shaded areas mark periods of a stable classification into either BW (red) or FW (blue) state; (7) topographical plots showing the mean EEG phase gradients (across 50 trials) corresponding to the classified BW and FW periods. C: Left: Response curve showing the probability of a FW state classification during a 3 sec DC stimulation as a function of input current. The dashed line marks the estimated threshold current (center of the sigmoid) at which the system switches wave states. Right: Temporal response functions for FW and BW responses to suprathreshold stimulation; the stimulus sequence used was a random step function with a fixed autocorrelation of 100 ms (∼ 1 cycle). Chance-level correlations (from random trial shufflings) are shown with 95% confidence intervals in the shaded area.

Feedforward projections between areas extending from supragranular layers (SG_E_) are inhibited through feedback from the subsequent higher area’s infragranular layers (IG_IB_). This connectivity constitutes a closed feedback loop that effectively reduces feedforward signals to the residuals between areas: we based this implementation on predictive coding principles which, as developed by Alamia & VanRullen (2019) with a much simpler approach, explains the emergence of traveling waves through feedback-loop connections. We demonstrate explicitly the relationship between our model and previous work in Supplemental Material (S1). The circuit described by this feedback loop, which carries the feedforward sensory input (as residuals or prediction-errors), will be denoted as the *FW-pathway*. We will discuss in greater detail how each pathway contributes to the oscillatory dynamics of the network below.

### Traveling wave dynamics in the cortex

We first investigated the behavior of the network to sensory stimulation, which we model by applying an external current to the lower end of the hierarchy, thus simulating a constant visual stimulus.

Panels (1) – (4) of Figure 1b summarize the network’s response during rest and to 1 second of direct current (DC) stimulation. In the absence of sensory stimulation, the intrinsically bursting nodes in the infragranular layers (IG_IB_) drive the network activity. The intrinsic rhythmic activity in these nodes synchronizes between areas due to their backward coupling in the network. The fixed inter-areal delay (ΔT = 12 ms) leads to a consistent phase-gradient corresponding to a backward (BW) traveling wave, propagating from higher to lower areas, which is visible in the map of mean-field activity (panel (4), showing averages per cortical area). At the onset of sensory stimulation, the rhythmic activity persists but switches to an opposite phase gradient, now corresponding to a forward (FW) wave, i.e., propagating from lower to higher areas. The activity traces (panels 2 and 3) show that the rhythmicity during stimulation is carried by both superficial and deep layers, in contrast to the baseline state in which superficial activity is largely unsynchronized. In a section below, we explore the extent to which the oscillation in the FW state is differentially determined by intrinsic IG_IB_ rhythmicity vs. the inter-areal feedback loop between supragranular nodes.

### Model dynamics at the scalp level

Next, we aimed at modeling the dynamics of traveling waves at the level of the scalp. To this end, we evaluated our model’s mean-field output by simulating an EEG signal using a standard forward model (with sources for Cx_1_, Cx_2,_ and Cx_3_ positioned bilaterally in occipital, parietal, and frontal regions, respectively). This additional step allowed us to quantify traveling wave direction and directly compare our simulations with experimental data. Panels 5 and 6 in Figure 1b summarize the results of this procedure for the same simulated trial shown in the panels above. The pattern of BW and FW waves remains visible in the spatiotemporal map of the simulated EEG (shown for electrodes on the mid-line, panel 5). We used an iterative plane fitting procedure (based on Zhang et al., 2018, see Methods) to obtain the one direction of propagation that best explains the spatial phase gradient for each time point (panel 6). The final classification into FW and BW states based on this value (shaded red and blue areas, with unshaded areas corresponding to states not classified as either FW or BW) matches the stimulus-dependent reversal of states that is visible in the mean-field activity (i.e., at the source level in the model, panel 4). Finally, the topography plots in panel 7 show the mean alpha-phase gradients (averaged across 50 model runs) corresponding to each state, limited to the electrode-ROI used for the fit, revealing a clear gradient of phases, reversed in the two states. In the following, we will primarily use the final classification output (i.e., FW or BW states) to explore the behavior of our model. It should be noted here that the two states are largely, but not strictly, complementary, as there may be cases where fits could not be classified into either category (i.e., the fit produced a correlation coefficient below a given null distribution, see Methods).

The single-trial example in Figure 1b already shows that the network’s wave state depends largely on the stimulus current injected, which is in line with experimental observations and previous modeling efforts (Alamia & VanRullen, 2019). To quantify the relationship between forward waves and visual stimulation, we mapped the probability of a FW state (across runs of 3 seconds of DC stimulations) to the stimulus amplitude, which revealed a sigmoidal response curve (Figure 1 c, left). This confirms that the network state depends systematically on the amplitude of the input current and allows the estimation of the threshold current at which the system switches wave states (at 0.79 nA in our simulations, dashed line).

Next, we aimed to characterize the network’s generalized impulse response, i.e., the model’s response to a brief (impulse) input. Specifically, we were interested in the time constants of the reversal response. To estimate this, we presented the model with suprathreshold random stimulation. Specifically, the stimuli were temporal sequences alternating randomly between ON- and OFF-stimulus periods, with a fixed stimulus current for the ON-periods (above the reversal threshold) and an imposed autocorrelation of 100ms (corresponding to approx. one cycle of the network’s oscillation, thereby the minimum resolution imposed by our wave fitting procedure). The resulting response functions for the classified wave states (obtained by cross-correlation with the stimulus sequence; Figure 1 c, right) exhibit a sharp response onset and a decay over a few oscillatory cycles, returning to chance-level around 700ms. This shape demonstrates two crucial properties of the model: 1) the reversal from the BW-to the FW-state is immediate, i.e., the first cycle after stimulus onset already propagates as a FW wave - notably, this mirrors the spatiotemporal pattern of physiological evoked responses to visual stimulation; and 2) the return to the baseline is comparably slow, indicating the network’s tendency to remain in the FW state for a few cycles once it has been evoked.

### Comparison of the model to human EEG data

After establishing the main properties of our model, we then compared its behavior to experimental observations, using EEG data published previously (Pang (庞兆阳) et al., 2020). In that study, human subjects were presented with a static luminance patch presented centrally in regular stimulus-ON/OFF sequences (of 5 seconds each). The EEG results showed a consistent increase of FW- and a decrease of BW-wave strengths during the stimulus-ON period (for a detailed report on the quantification of waves in that analysis, cf. their Methods). We first replicated this pattern of findings by applying the same phase-gradient analysis that we use to evaluate our model output (cf. Figure 1b, panels 6 & 7, and Methods). The left panel in Figure 2a shows the mean probability of observing a stable FW or BW state over time across trials and subjects. Here and in all other panels, the grey shaded area represents the time-period the stimulus is on. As shown in the right panel of Figure 2a, our model captures the empirical response pattern well. Note that the aim of our simulations was not to reproduce the signal-to-noise (SNR) ratio of the actual EEG data, and the difference in scaling for both classification probabilities (FW and BW) is an expected consequence of such a difference in SNR. As expected from the impulse response function, our network exhibits the same reversal from a BW wave state during baseline to a FW wave state upon stimulus onset, as present in the experimental data. The transition times for stimulus on- and offset were comparable between experimental and simulated data, accounting for a fixed processing delay in the physiological data, possibly due to processing in the retina and subcortical nuclei (time to first peak at stimulus onset - data: 363 ms, model: 61 ms; return to baseline (corrected for linear slope) - data: 744 ms, model: 441 ms). Thus, in summary, our model can qualitatively reproduce the population-average dynamics of the traveling wave response to visual stimulation.

**Figure 2.**
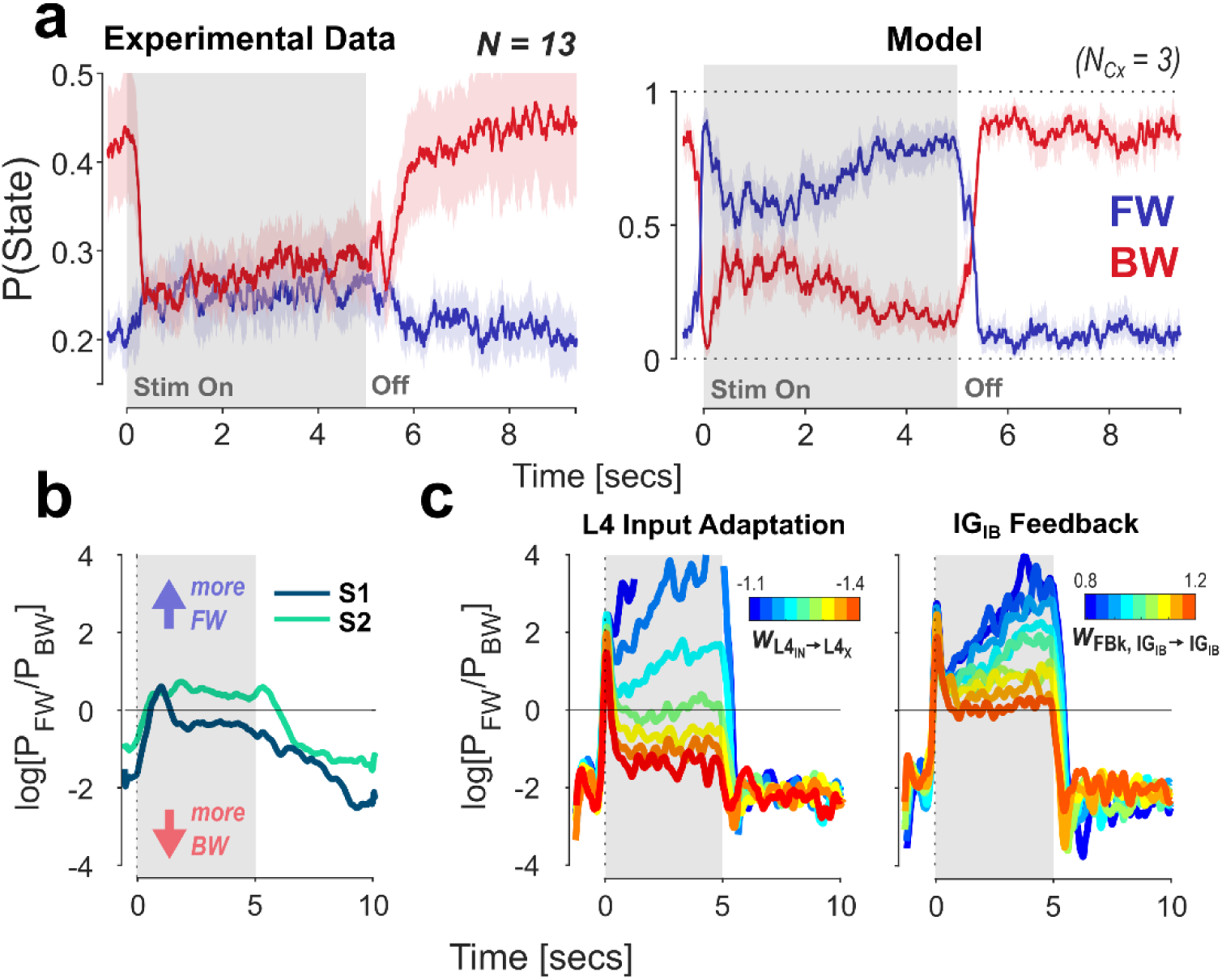
Comparison of our model to experimental EEG data (from Pang (庞兆阳) et al., 2020). The stimulation consists of a 5 sec static input (shaded area) followed by a rest period. A: Left: Grand average time-courses of mean FW / BW probabilities from the fit on the experimental data. Shaded areas represent +/-1 SD to indicate between-subject variability. Right: Probability time-courses obtained from 100 model runs (trials). Here, shaded areas represent the 95% CI based on trial-variability. Stimulus amplitude was 0.79 nA (threshold value obtained from the response curve, cf. Figure 1 c). B: Experimental data: responses from two subjects, shown as the log-ratio between wave states over time. Variability in the response shape is visible in the baseline offset and the amount of response adaptation after the initial peak. C: We explore two parameter variations to model the variability observed between subjects. FW/BW ratio are shown as a function of different parameter values. Left panel: the weight of inhibitory recurrent feedback on the input in Layer 4 (w_L4IN_→_L4X_); Right panel: the weight of excitatory feedback between infragranular pacemakers (w_IGIB_→_IGIB_). All weights were varied together for all areas in the network.

However, a closer analysis of the data from Pang et al. reveals considerable inter-individual differences in the responses. Figure 2b shows the response time courses for two subjects, represented here as the log ratio of FW and BW state probabilities. These illustrate the two primary sources of variability in the data we consider relevant for our model: 1) the baseline ratio between the two states (i.e., the individual strength of the bias towards BW waves in the absence of stimulation), and 2) the temporal evolution, i.e. shape of the response within the 5 sec of the stimulus-ON state. We turn to the former in more detail below, where we show that the extension of the cortical model by a pathway through the pulvinar reproduces such bias in the network resting state. Concerning the variability in our model’s response shape, we explore its dependence on two key parameters. The first is the amount of suppression exerted on the feedforward signal through neural adaptation at each area. In our model, adaptation is achieved through recurrent coupling to a local inhibitory population in Layer 4 (i.e., L4_IN_ in Figure 1A). This allows the strength of input adaptation to be varied directly through the weight of the inhibitory connection. The left panel in Figure 2c shows the effect of varying this parameter on the network response (expressed as the log-FW/BW ratio, as in panel b), given the same stimulus pattern presented before (i.e., 5 seconds of DC stimulation). As expected, the evoked FW response is increasingly suppressed with stronger adaptation, while the first cycle of the response remains unaffected. The dynamics of this rapid decay follow directly from the time constant of the Layer 4 recurrent inhibition (neural integration time + synaptic time constants).

The second parameter we varied is the strength of the excitatory infragranular feedback connection (IG_IB_ in Cx_i_ to IG_IB_ in Cx_i-1_) to determine whether the network’s wave state can also be biased through modulation of the BW pathway (higher weights correspond to stronger feedback). The results are shown in the right panel in Figure 2 c. The synaptic connection between IG_IB_ nodes is excitatory, thus higher weights correspond to stronger feedback. The effect of feedback weight largely mirrors the one obtained for the input adaptation strength, with stronger feedback leading to a weakened sustained FW response. This result illustrates the continuous competition between the FW and BW pathways, which will be discussed below.

These results demonstrate that the response dynamics can already be flexibly modulated through the variation of two synaptic weights in the network, possibly explaining some inter-subject variability in EEG recordings. Naturally, other parameters may play a role in modulating the pattern and strength of traveling waves, such as the weight of the feedforward connection (SG_E_ → L4_X_), or the di-synaptic inhibitory feedback to superficial layers (IG_IB_ → SG_IN_ → SG_E_). Besides explaining inter-subject variability, differences in specific parameters between cortical areas may be instrumental for the modeling of specific functional networks, e.g., a systematic increase in neural timescales along cortical hierarchies to match empirical observations (Gauthier et al., 2012; Murray et al., 2014), or in formulating specific predictions about alteration in traveling wave patterns in clinical populations (Alamia et al., 2024).

### Subcomponents of the network: FW and BW pathways

Next, we investigate the mechanisms that drive the generation and propagation of FW and BW waves in our model. As described above, the network architecture can be separated into two components: 1) the BW-pathway, a set of backward coupled nodes with intrinsic rhythmicity in the infragranular layers (IG_IB_), and 2) the FW-pathway, an inter-areal closed loop in which signals are passed forward from superficial layers into Layer 4 (SG_E_ → L4_X_), then locally through superficial to deep layers, and finally as feedback into superficial layers of the lower area (IG_IB_ → SG_IN_). This implementation is inspired by previous modeling and theoretical work considering a predictive coding perspective (Alamia & VanRullen 2019, Bastos et al. 2012). In the following, we will examine how the rhythmicity of the network (specifically, its dominant temporal frequency) is influenced by the mechanisms generating oscillations in either pathway. Figure 3a shows the distribution of spectral power in supra- and infragranular nodes at baseline (gray lines) and during sensory stimulation (colored). As illustrated by the single-trial example in Figure 1b, the intrinsically bursting property of the IG_IB_ nodes leads to a constant stable rhythm in the infragranular layers that increases in frequency only slightly during stimulation (see below). In contrast, rhythmic SG_X_ activity only emerges during stimulation, matching the frequency of the IG_IB_ pacemaker. Mapping the peak frequency of activity in both compartments against the input current (Figure 3b) confirms this pattern. Figure 3b also reveals that the two frequencies are tightly coupled, showing a slight increase in frequency in both IG_IB_ and SG_X_ following an increase in input current above threshold (0.83 Hz and 1.32 Hz increase over the suprathreshold range for IG_IB_ and SG_X_, respectively). To disentangle the influences of the two pathways on the network’s temporal frequency, we leverage the fact that the mechanisms generating oscillatory activities within each pathway are different. While rhythmicity in the BW pathway is a property of the nodes and thus independent of connectivity, it emerges through a feedback loop in the FW-pathway. In the latter, the oscillation frequency depends (in addition to neural integration times), crucially, on the inter-areal delay (for a detailed derivation of this, we refer the reader to Alamia & VanRullen, 2019). Building on this dependency specific for the FW, but not the BW pathway, we use a systematic variation of the inter-areal delay to disentangle the two pathways during visual stimulation. To identify the interaction between the two pathways, we compare the changes in the dynamic between two different model configurations while varying this parameter (as summarized in Figure 3c, d). In the first configuration, we test the full model with both pathways in place (left panels in c and solid lines in d). In the second configuration, the FW-pathway has been isolated by removing both the intrinsic rhythmicity and the inter-areal feedback in the BW-pathway (right panels and dashed line). As predicted, in both model versions, increasing the inter-areal delay caused a decrease in the dominant frequency of SG_X_ activity (Figure 3c, lower panels). In the full model, this decrease leads to a divergence between SG_X_ and IG_IB_ peak frequencies (panel d) for delays larger than approximately 18 ms. In this diverging regime, oscillatory *power* in the SG_X_ nodes also decreases due to the loss of synchronization between the two pathways. This desynchronization removes the IG pacemakers’ facilitating influence on the FW-pathway activity. For larger delays, SG_X_ oscillatory power thus returns to the same level it can sustain without the BW pathway, i.e., in the FW-isolated configuration (panel c, note the difference in scales between colormaps). These two observations (i.e., the divergence in temporal frequency and the loss of oscillatory power) suggest at least some degree of independence between the two pathways, even when they are interconnected in the full model. Importantly, however, the intrinsic frequency of the isolated FW-pathway is consistently lower than that observed in the full model (SG_X_ peak frequency difference at ΔT = 12ms: 2.81 Hz). This is true for all tested delays, i.e., both in the non-diverging and the diverging regime. This offset in frequency directly shows that the intrinsic rhythmicity of the IG_IB_ pacemakers largely dominates the frequency of the FW pathway. It is important to note that all of the results shown in Figure 3c & d represent oscillatory activity during the FW wave state. Thus, even though the *temporal frequency* of the network follows the infragranular pacemaker (i.e., the BW pathway) and is more or less stable between stimulus ON and OFF states, the *spatial propagation* of the activity (i.e., the relative phase between areas) reverses when the FW pathway is activated. In sum, we showed here that the two pathways have independent mechanisms to generate oscillations. However, in the fully connected model and for a fixed delay of 12 ms, the infragranular pacemaker largely determines the temporal frequency of the network. In contrast, the involvement of the FW pathway only determines the directionality of the phase gradient.

**Figure 3.**
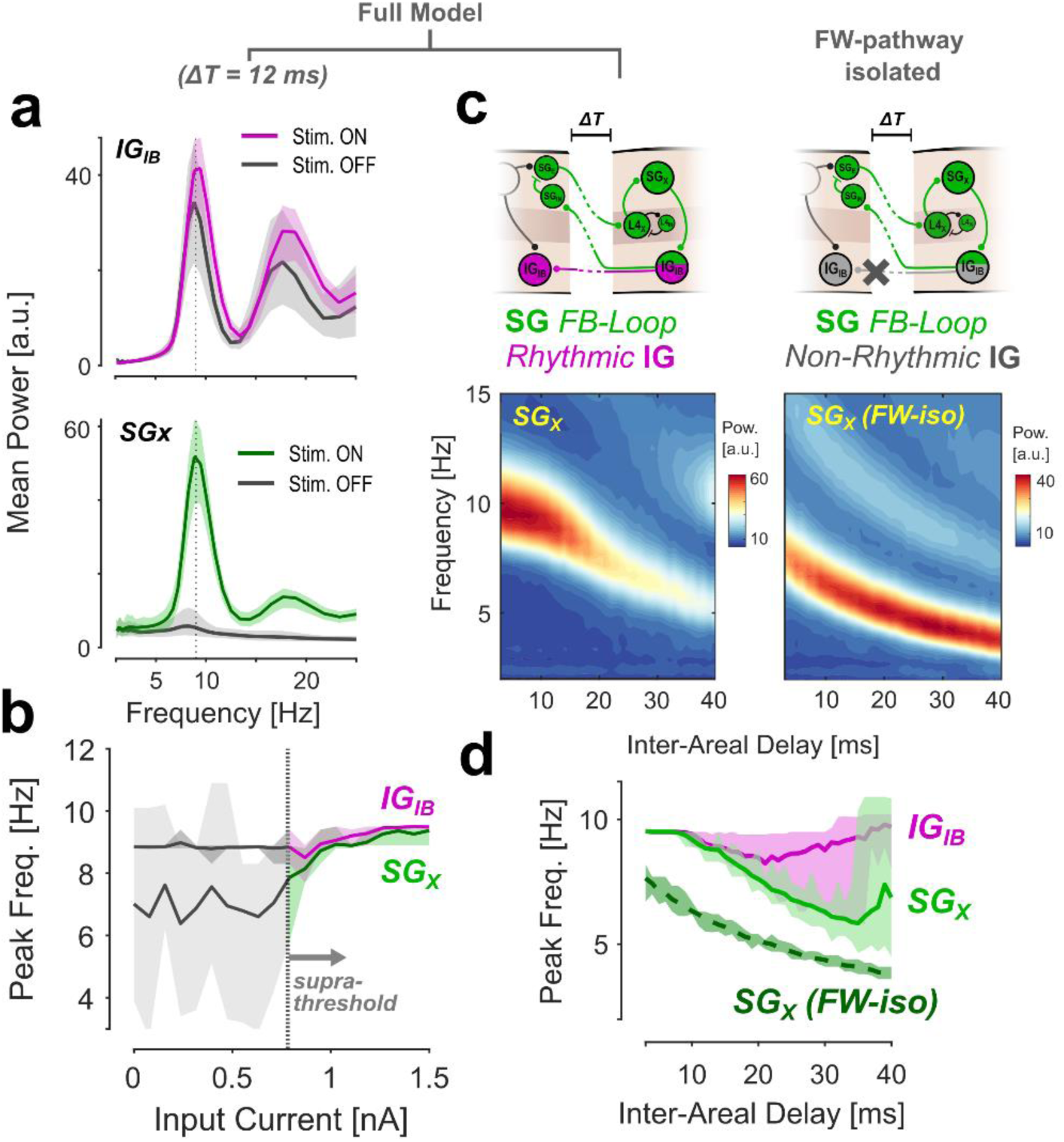
Isolation of FW- and BW-pathways in the model. A: Mean power spectra of all IGIB (_to_p) and SGX activity (bottom) during stimulus On and Off periods. B: Peak frequency for both node types as a function of input current. The two frequencies are tightly coupled for stimulation above the threshold. C: Variation of inter-areal delay in two configurations. The left panels show the full model; in the right panels, the FW pathway has been isolated by reducing the infragranular nodes to a passive relay (without intrinsic rhythmicity). The bottom panels show the mea_n_ power spectra for SG_X_ as a function of delay. D: Peak frequencies for the data presented in C. The two pathways diverge for larger delays in the full model, while the intrinsic frequency of the FW pathway is consistently lower. Shaded areas in A, B & D show 95% quantiles for the variability between model runs.

### The pulvinar pathway as a mechanism to bias traveling wave direction in the cortex

After demonstrating that the proposed cortical model can explain the dynamics of alpha traveling waves at the EEG level, we next investigate the role of the cortico-thalamic network, specifically the pulvinar, to explore how its connectivity affects wave dynamics in the cortex. The cortico-pulvinar circuitry (including the thalamic reticular nucleus, TRN) comprises a complex set of connections involved in seemingly diverse functional roles (for overviews, see, e.g., Shipp, 2003; Saalmann & Kastner, 2011; Sherman, 2016; Grieve et al., 2000; Kaas & Lyon, 2007). A detailed analysis of how each part of that circuitry could modulate wave dynamics is beyond the scope of this study. Here, we focus on the main Cx-Pul-Cx pathway that connects any two areas connected directly in the cortex (i.e., replication principle, Shipp, 2003). Furthermore, within this pathway, we focus on the driving feedforward projections extending from deep layers to the pulvinar and then to Layer 4 of the higher cortical area (Sherman & Guillery, 2002) (while noting that both feedback, as well as non-driving, modulator projections also exist between some cortical regions and the pulvinar; see Discussion). Importantly, we also do not consider the cortico-thalamic system as an additional, independent generator of alpha oscillations, as suggested by some experimental findings (Saalmann et al., 2012; Zhou et al., 2016) and modeling (Cortes et al., 2021). As before, we aim to keep our model as simple and general as possible by reducing connectivity to what is likely shared between most hierarchically connected visual areas.

Figure 4a shows the schematic of our model extended by the trans-pulvinar pathway (in orange). In the logic of the FW- and BW-pathways described above, the connection through the pulvinar may be seen as running parallel to the FW-pathway in that it provides a second feedforward connection into Layer 4 of each area. The critical distinctions from the FW-pathway are 1) that it is unidirectional, i.e., it does not support the generation of its own oscillation through a feedback loop, and 2) that it is active in the absence of external input by the intrinsic activity propagating from the infragranular nodes. Thus, the pulvinar in this configuration provides an excitatory shortcut feeding spontaneous IG_IB_ activity into the FW pathway.

**Figure 4.**
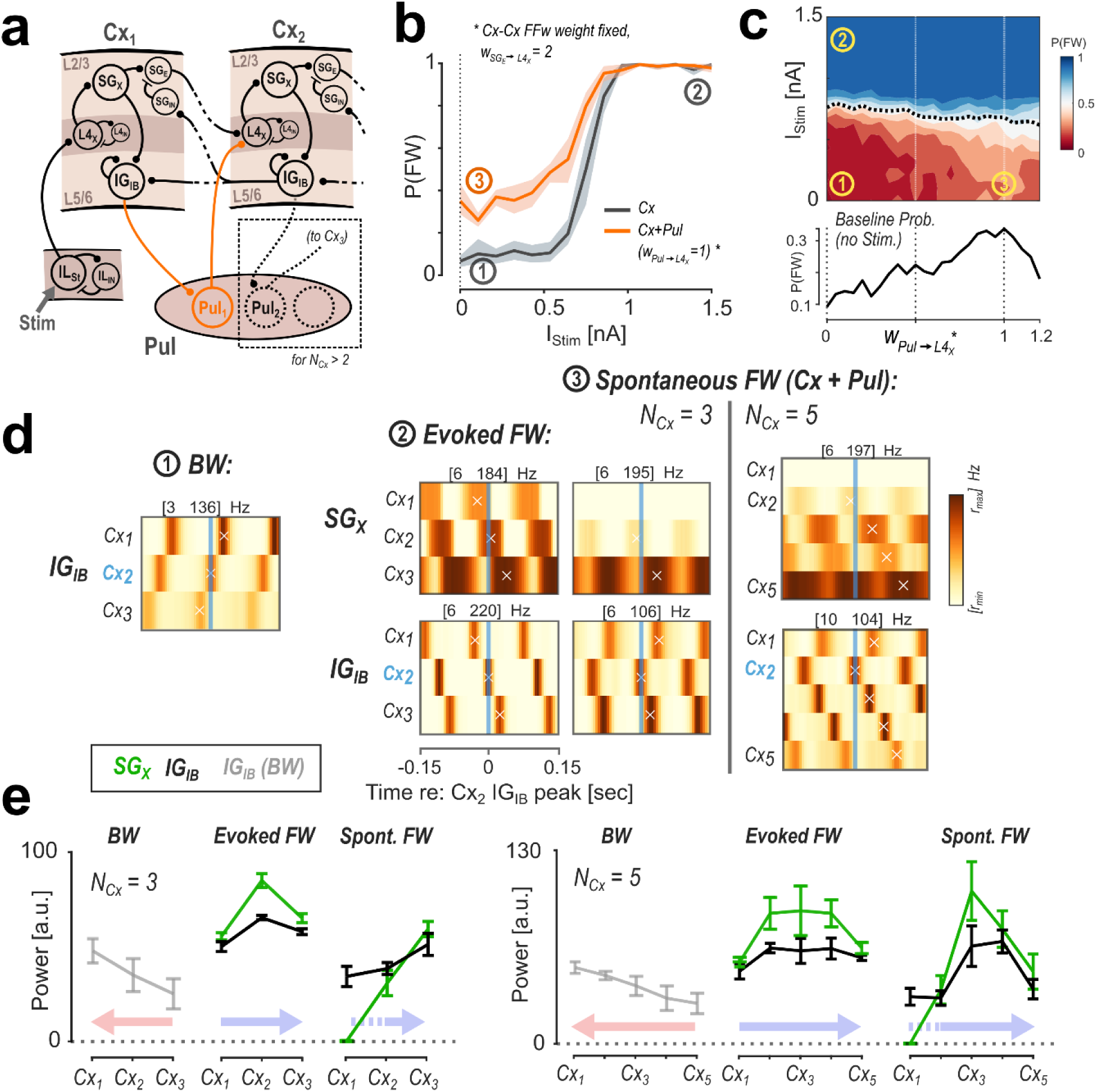
Modulation of FW waves by the pulvinar. A: Model architecture with the added pulvinar pathway. For every two successive areas, one pulvinar node relays a feed-forward connection from infragranular layers into Layer 4. All cortical connectivity remains the same as before. B: Response curves for the Cx-only (grey, same as Figure 1 c) and Cx+Pul (orange) configurations. C: Full map of FW state probability as a function of pulvinar engagement (w_Pul_→_L4X_). The dashed line indicates the estimated response threshold. The bottom panel shows the baseline probability at I_Stim_ = 0 nA. D: Spatiotemporal maps of cortical activity associated with each wave state labeled 1-3 in B & C. Each sub-pan_el_ sho_ws_ SG_X_ / IG_IB_ activity across cortical areas and time, averaged over epochs centered on the p_ea_k activity at Cx_2_-IG_IB_. For spontaneous FW waves, results for a 5-area version of the model are shown for comparison. The maps show that the spontaneous FW waves associated with pulvinar engagement originate in Cx_2_, i.e. are shifted downstream compared to evoked FW waves. E: Distributions of mean oscillatory power across the network for 3-area (left) and 5-area versions, comparing the wave states described above. Error bars represent +/-1 SD between model runs. Colored arrows show the direction of propagation.

Figure 4b shows how the addition of this pathway affects the scalp-level wave dynamics in our model: for a high pulvino-cortical weight, the response curve for the FW wave state (same as Figure 1c) is shifted to a higher baseline probability at the lower bound (while the response threshold, i.e., the sigmoid inflection point, remains similar). In other words, spontaneous network activity (without any input current) is more likely to generate FW waves when the pulvinar pathway is engaged. This pattern is also visible when continuously mapping the response curve to the weight of the pulvino-cortical connection (Figure 4c). The dashed line in the upper panel marks the estimated threshold input current (50% point of the sigmoid), while the lower panel shows the baseline probability. The decrease of the baseline probability for weights > 1 here is explained by an overstimulation of the downstream cortical areas due to the increasing net current injected into these areas via the pulvinar connection (note that the cortico-cortical FFw weights are fixed at w = 2).

Since they are based on scalp-level classification, the response curves in panels b & c do not allow conclusions about the activity in the cortex that underlies the FW-biased state. To elucidate this, we compared three states (marked in panels b & c): 1) the stable BW wave state (without external input) in the cortical-only model, i.e., with the pulvinar disconnected, 2) the stable, stimulus-evoked FW state in the same configuration (cortex-only), and 3) the state without external input in which spontaneous FW waves emerge as a result of the added pulvinar pathway (w_Pul_→_L4X_ = 1). For each state, we consider the spatial activity maps (obtained by concatenating firing rates from nodes of the same type across the cortical hierarchy) in SG_X_ and IG_IB_ nodes after averaging across epochs centered on the peaks in the rhythmic IG_IB_ activity at Cx_2_ (shown in Figure 4d). We chose this reference for its position in the center of the network (Cx_2_ for N_Cx_ = 3) and because the IG_IB_ node is at the intersection between FW and BW pathways. The maps for BW (1) and evoked-FW states (2) show the expected pattern of propagation with Cx_3_ and Cx_1_ activity leading in both laminar compartments, respectively (Figure 4 d). In contrast, during the spontaneous-FW state (3), activity at Cx_2_ leads the other areas, while the supragranular nodes in Cx_1_ remain inactive. This result illustrates how the spontaneous FW waves are elicited in the Cx + Pul configuration: spontaneous IG_IB_ activity in Cx_1_ is relayed to Cx_2_ via the pulvinar, activating the FW pathway in the same way as the external input does in Cx_1_. This results in a spontaneous FW wave spatially originating at Cx_2._ The infragranular activity in Cx_1_ disrupts the spatial phase gradient, leading to an unstable classification at the scalp level.

The activity pattern of spontaneous FW waves becomes more evident when considering a network with more than three areas. The two maps on the right in panel d show the case for N_Cx_ = 5 as an example. Here, the consistency in the FW wave is more clearly visible, while the leading activity remains at Cx_2._

Lastly, the two FW wave states (evoked/spontaneous) also differ in their spatial distribution of activity levels and oscillatory power (Figure 4e). As a result of cascading effects in the FW- and BW-pathways, overall activity levels in either state naturally increase with the direction of propagation (i.e., more oscillatory power at regions trailing in phase; panel d; BW: *IG_IB_*, FW: *SG_X_*). Since physiological firing rates are subject to strong normalization from various sources (and were not the target of our modeling), differences in the baseline firing rate can be disregarded. Instead, we tentatively interpret oscillatory power as a measure for the spatial distribution of wave amplitude (shown in Figure 4e) (note that we deliberately refrain from modeling power topographies at the scalp level as these depend significantly on assumptions about the size of active populations in each area). The power distribution in the evoked FW state peaks at the center of the network for both versions of the model (3-/5-stage). For spontaneous FW waves, this distribution is skewed towards downstream areas due to the shift in wave origin from Cx_1_ to Cx_2_.

In summary, adding the pulvinar feedforward pathway in our model leads to the emergence of FW wave patterns in the baseline state while leaving the response to external stimulation unaltered. These spontaneous FW waves differ from those evoked by stimulation in that their spatial origin is shifted to the first area downstream of pulvinar feedforward activation. In our simulations, where the pulvino-cortical weights w_Pul_→_L4X_ are identical across regions, the leading area is the second lowest (i.e., Cx_2_), but a differential modulation of these weights would allow for a flexible control of the spatial distribution of the wave.

### Lateralization of cortical wave states

In the previous section, we demonstrated that our network’s wave state (FW/BW) can be biased through the pulvinar pathway. Interestingly, a possible cognitive function associated with this modulation type is the control of attentional allocation. Empirical observations have suggested that traveling wave direction at the scalp level is lateralized as a function of visual attention (Alamia et al., 2023), similar to alpha-band oscillations (e.g., Händel et al., 2011; Sauseng et al., 2005; Thut et al., 2006). As we will demonstrate in the following, our model can reproduce such lateralization of wave states consistent with hemifield-directed attention through modulation of pulvinar engagement. Specifically, we adapted our simulations and wave fitting procedure to differentiate between hemispheres (Figure 5a). The novel network consists of two independent thalamo-cortical streams, each following the same architecture and connectivity as above. In this configuration, we did not include direct cortical or subcortical interconnections between the two hemispheres. Each hemisphere receives its input current into the first area, simulating left and right visual field stimulation. After projecting all cortical sources together to the EEG, we estimate wave states in two separate regions of interest (ROIs) lateralized to either hemisphere in the sensor space (Figure 5a). We first tested the effect of lateralization in this model by passing the input current to only one hemisphere. Figure 5b shows the probability time courses for FW state classification in both ROIs across a 1 sec left-hemisphere DC pulse stimulation. The response clearly shows the expected lateralization to the stimulated side. Notably, the opposite ROI does not show a residual response (an intuitive expectation one may have from the spatial mixing of signals in the EEG projection) because the right hemisphere remains in a stable BW state during stimulation. This demonstrates, as a pre-requisite for the modeling of hemispheric modulation, that simultaneous waves traveling in opposite directions between the two hemispheres can be picked up reliably from a single simulated EEG signal. However, the effect of signal mixing is evident in an overall reduction of classification consistency (here, I_Stim_ = 1nA, compare with Figure 1c). For reference, the topographical plot on the right shows the lateralization of the same signal in the time-domain (root-mean-square (RMS) within the first 100 ms after stimulus onset). Lastly, after establishing that our model can generate lateralized wave states (simultaneous opposing waves), we also investigated whether it can reproduce hemispheric modulation of waves traveling in the same direction. For this, we mapped responses in the two ROIs separately while passing the same shared input current to both hemispheres. Importantly, we varied the weight from pulvinar to cortex selectively for the left hemisphere, while keeping the weight for the right fixed at a low initial value (Figure 5c). The results reveal that the response curve for the left ROI is selectively modulated, confirming the shift in baseline of the FW state probability with increased pulvino-cortical weight. Additionally, due to the lower SNR in the responses (as compared to the non-lateralized version), the modulation persists even in the supra-threshold state (high input current), in which a FW wave is evoked in both hemispheres. This simulates the evaluation of EEG recordings better than the more ideal response curves in Figure 4b. Overall, our results show that a lateralized modulation of pulvino-cortical weights can lead to a hemispheric bias in FW wave consistency, similar to those obtained for selective visual attention. As shown experimentally (Alamia et al., 2023), this lateralization can be reliably detected at the scalp-level, despite mixing signals from both hemispheres in the EEG.

**Figure 5.**
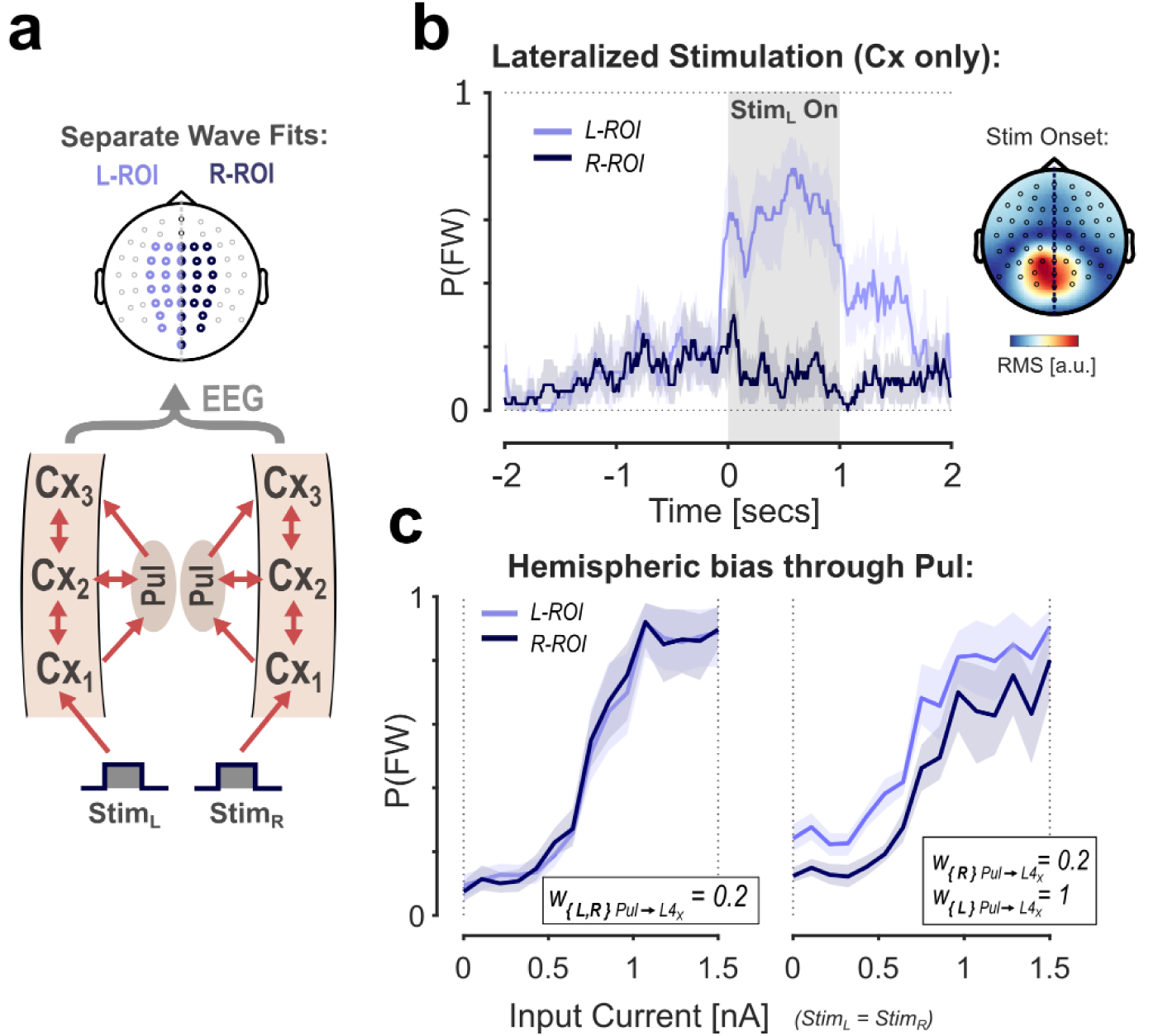
Lateralization of cortical waves and hemispheric modulation by the pulvinar. A: Illustration of the multi-scale model with a separate cortical stream per hemisphere. Waves states are estimated from separate ROIs based on a single EEG projection. B: FW state probabilities for the two ROIs in response to a 1 sec left hemisphere stimulation. The responses clearly show the expected lateralization of wave states. The topographical plot shows the root-mean-square (RMS) within the first 100 ms after stimulus onset to illustrate the lateralization in the time-domain. C: Response curves for each ROI with equal (left panel) and left-lateralized (right panel) pulvinar engagement (w_Pul_→_L4X_). The response shift shows that the effect of hemispheric bias from pulvinar modulation can be measured at the scalp level.

## Discussion

We constructed a multiscale network model that reproduces the pattern of alpha-band travelling waves at the level of the scalp, building on physiologically plausible neural dynamics. The modulation of FW/BW propagation along the cortical hierarchy by stimulus input qualitatively matches that observed in human EEG data. We have demonstrated that the generation of FW and BW waves in our model relies on distinct sub-components (FW- and BW-pathway, respectively). Lastly, we have further shown that the addition of a redundant feed-forward pathway through the pulvinar biases the network’s dynamics towards the FW-wave state in the resting (stimulus OFF) state. This type of control can be used to simulate hemispheric modulation of traveling wave direction, as measured via EEG.

### From source to sensor space: Linking scalp-level traveling waves to laminar circuits

A number of EEG studies in recent years have found evidence that traveling waves at the scalp-level are linked to cognitive and sensory functions (Alamia et al., 2023; Alexander et al., 2008, 2009; Ito et al., 2005; Pang (庞兆阳) et al., 2020; Patten et al., 2012). Similarly, traveling waves at different scales have been observed in a variety of paradigms, both intracortically and in recordings from the cortical surface (Aggarwal et al., 2022; Davis et al., 2020; Zabeh et al., 2023; Zhang et al., 2018). It remains unclear how the findings at these two levels (scalp and cortical) relate to each other. The source-level correlates of the scalp waves are ambiguous. This is in part due to the well-established forward problem (for which standardized solutions exist, see Hallez et al., 2007). However, more importantly, phase gradients at the scalp can equally result from intracortical travelling waves and from sequential activation of phase-lagged stationary sources (for different theoretical accounts, see, e.g., Hindriks et al., 2014; Zhigalov & Jensen, 2023). Lastly, establishing a direct link with (simultaneously recorded) cortical activity is methodologically challenging.

Our multiscale model constitutes a first effort towards bridging this gap. It describes a biologically plausible laminar circuit to explain the traveling wave dynamics observed at the scalp-level using EEG recordings. It also generates predictions about the cortical activity expected during FW and BW wave states, which could be tested experimentally in future studies combining multi-scale measurements.

A central prediction made by our model is that the scalp-level propagation of waves corresponds to a propagation along the hierarchy, i.e., a successive activation of cortical areas. This is consistent with the conclusion drawn by other authors that have identified long-range cortico-cortical fibers as the likely substrate of waves measured by the EEG based on conduction velocities (Muller et al., 2018; Nunez et al., 2001; see also the dependence of wave stability on interareal delays in Alamia & VanRullen, 2019). Notably, this is not equivalent to a continuous wave in the cortex, as our model does not make predictions about cortical tissue between two areas in the hierarchy, or non-activated regions within an area. Yet, a continuous propagation seems more likely, given experimental evidence of continuous traveling waves at meso- and macro-scales (Davis et al., 2020; Zhang et al., 2018). However, several different mechanisms of propagation would likely interact to generate a continuous wave from the successive activation of areas predicted by our model (see Muller et al., 2018, for an overview of the neural mechanisms driving propagation at different scales). Additionally, a recent study showed that scalp-level waves elicited during visual stimulation can sufficiently be explained by just two cortical sources (Zhigalov & Jensen, 2023). Our model is in line with this in principle (given that it generalizes to all cases with more than one area), however, our hypothesis would be that the number of sources needed to explain the wave depend on the particular functional pathway activated. This could readily be tested in future EEG/MEG experiments. Another important prediction from our model is that FW and BW wave states are associated with different laminar distributions of alpha activity. Specifically, alpha power in superficial layers should be stronger during the FW state (cf. Figure 3 a), whereas rhythmic activity in the deep layers is continuous (and only switches in directionality of the phase offset between states). While a dependence on global wave state remains to be tested experimentally, there is already a large body of evidence showing that different neural rhythms have distinct laminar distributions. Generally speaking, low-frequency activity (including alpha) seems to be stronger in infragranular layers, while higher frequencies are more prominent in supragranular layers (Lopes Da Silva & Storm Van Leeuwen, 1977; Maier et al., 2011; Mendoza-Halliday et al., 2024). Pacemaker neurons in Layer 5 have been identified as a potential generator of the infragranular alpha (Steriade et al., 1990; Sun & Dan, 2009), and the main generator for alpha in our model (IG_IB_) reflects these findings. However, other studies report evidence for separate generators in both deep and superficial layers (Bollimunta et al., 2008, 2011; Van Kerkoerle et al., 2014). In line with this, the network architecture in our model allows for the generation of a second alpha rhythm through interareal feedback to supragranular layers. It will be crucial for our understanding of cortical traveling waves to investigate how potential different alpha generators interact across different cortical laminae.

### Alpha traveling waves as correlates of forward- and backward-directed communication in the cortex

Our model distinguishes largely between two global states, with waves propagating either FW or BW along the cortical hierarchy. Here, these two states simply reflect the directionality of the phase gradient between areas, but there is broad evidence to support a relationship between the direction of propagation and cortical communication. For example, in EEG recordings, alpha-band FW waves have been linked to active visual processing and attention, whereas alpha-band BW waves are more prominent during rest and attentional suppression (Alamia et al., 2023; Pang (庞兆阳) et al., 2020).

In contrast to this, in recent years intracortical recordings have led to the emerging view that communication directions are separated among different frequency ‘channels’, with higher frequencies (gamma) carrying feedforward and lower frequencies (alpha/beta) carrying feedback signals (A. M. Bastos et al., 2015; Jensen et al., 2015; Michalareas et al., 2016). The two channels interact in that gamma activity (in supragranular layers) is modulated by the phase of (infragranular) alpha (Spaak et al., 2012), in line with the concept of cross-frequency coupling (Canolty & Knight, 2010).

Our model, supported by experimental data (Alamia et al., 2023; Pang (庞兆阳) et al., 2020, Alamia & VanRullen, 2019), challenges the notion that alpha rhythms represent purely feedback signals. Instead, it suggests that alpha provides a constant global rhythm whose phase gradient can switch directionality to support communication in either direction. Importantly, the high-frequency (gamma) channel could be easily incorporated into our model, e.g. by a separate node in supragranular layers generating gamma through a pyramidal-interneuron gamma (PING) mechanism. The supragranular activity in the current model would translate to an amplitude modulation of the gamma envelope in that node. In this implementation, the alpha phase modulating the gamma activity would be generated by the inter-areal feedback connection. Additionally, a second, direct projection to the supgranular layers from deep layers of the same area is also likely (Dantzker & Callaway, 2000; Xu & Callaway, 2009). While the cross-frequency interaction was not the target of our modeling, we consider this integrated model (i.e., one with seprate alpha- and gamma-frequency channels) here because the distinction between alpha-phase-*modulated*- and *intrinsic* alpha activity may explain the seemingly contradicting roles of alpha between EEG and laminar recordings. In other words, the scalp-level alpha during the FW wave state may represent rather the phase-modulated gamma activity in supragranular layers, while BW alpha would represent infragranular intrinsic alpha. Previous studies provide some support for this more flexible role for alpha oscillations (as reflecting cortical communication more generally, as opposed to a feedback-specific channel). Using ECoG in humans, Bahramisharif et al. (2013) found that the amplitude of gamma activity is modulated by the local phase of large-scale alpha traveling waves, effectively resulting in the spatial propagation of gamma bursts. Chapeton et al. (2019), without characterizing traveling waves specifically, showed experimentally that the communication between regions is optimal when their alpha activity is coherent and phases are aligned to match the conduction delays between them. A similar framework of alpha-gated communication has been proposed by Bonnefond et al. (2017), and demonstrated in a biologically plausible network model (Quax et al., 2017). Lastly, Halgren et al. (2019), using ECoG and sEEG in humans, found traveling alpha waves that propagate towards the occipital pole during rest, consistent with our prediction that the resting alpha rhythm is backward-directed. They also report evidence of a reversal of waves to the FW direction after opening of the eyes in a macaque (but only in some of their human subjects, see their Supplemental Figure S4). Future studies could establish the dynamics of traveling wave direction during similar transitions from rest to active states.

Interestingly, Halgren et al. localize the laminar source of alpha selectively in the superficial layers, contrary to previous studies and our model predictions that assume either distributed or infragranular sources. It is possible that the discrepancy in the literature partly reflect the use of different analytical measures (current source density (CSD) in Halgren et al. vs. local-field potential (LFP) in most studies). However, it seems possible that waves that are generated by infragranular pacemakers down-stream in the visual hierarchy propagate further via short-range connections in superficial layers. This would be an indication that different laminar distributions are to be expected at the presumed source of a wave and at remote cortical regions through which it propagates.

### A Predictive coding interpretation of the FW and BW states

The interlaminar and interareal connectivity in our model is inspired by the hierarchical predictive coding model developed by Alamia & VanRullen (2019). It also follows the microcircuitry proposed by existing laminar models of predictive coding (A. M. Bastos et al., 2012; Shipp et al., 2013; Shipp, 2016). Thus, even though our model itself is dynamical and does not include a feature space, it supports the computation and passing of prediction errors between areas. Specifically, in line with previous studies, it predicts that this error computation occurs in superficial layers. This is further supported by recent electrophysiological studies who found evidence for a neural comparison between top-down and bottom-up signals in Layers 2/3 (G. Bastos et al., 2023; Gallimore et al., 2023; Hamm et al., 2021; Jordan & Keller, 2020). Complementary to this, Thomas et al. (2024) were able to show with 7T fMRI in humans that stimuli with large prediction errors were represented selectively in superficial layers.

While our model is thus well in line with experimental evidence supporting predictive coding, the specific functional role of alpha traveling waves in these computations remains unclear. In particular, considering the retinotopic representation at early stages of the visual hierarchy (V1/V2), it needs to be shown to what extent the reversal of wave direction is spatially selective, to support the feedback of spatially informative predictions to these areas. There is already some evidence from the EEG that different wave states can co-occur, triggered by stimuli at separate positions in the visual field (Alamia et al., 2023; Lozano-Soldevilla & VanRullen, 2019), and our own simulations with two separated hemispheric streams (Figure 5) also demonstrate this. However, the low spatial resolution of the EEG limits the conclusions to be drawn. Future studies could expand our dynamical model to include a feature space, in order to investigate how feature-based predictions (such as about the position of a stimulus) interact with traveling wave dynamics.

Interestingly, a recent computational model showed that meso-scale waves, traveling horizontally across retinotopic space within an area, can also carry predictions about future visual input (Benigno et al., 2023). This corroborates experimental evidence that similar meso-scale waves occur spontaneously in the cortex and modulate local spiking activity as well as perception (Davis et al., 2020). An interesting venue for future studies will explore how the two types of waves (meso-scale within areas vs. feedforward/feedback between areas) interact, both dynamically and in the potential encoding of predictions and prediction errors.

### The role of the pulvinar

Our simulation results showed that the addition of second pathway through the pulvinar biases our network towards the FW wave state. A detailed analysis revealed that the spontaneous FW waves at rest propagated from the second cortical area (the first receiving input from the pulvinar), in contrast to stimulus-evoked FW waves, which progapated from the lowest area.

These results are in line with a number of studies (experimental and modeling) showing that the pulvinar plays a key role in the control of communication between cortical areas (Fiebelkorn & Kastner, 2019; Jaramillo et al., 2019; Quax et al., 2017; Saalmann et al., 2012; Zhou et al., 2016). In particular, the modeling effort by Jaramillo et al. (2019) links the pulvinar to diverse cognitive functions as a flexible gating mechanism in the flow of information between cortical areas. Together with our results, this opens an intriguing connection between the modulation of traveling wave direction and cognitive functions. There is initial evidence for a similar link in the hemispheric modulation of wave direction with spatial attention (Alamia et al., 2023). The decision-making paradigm modeled in Jaramillo et al. may provide another venue in this direction for future EEG studies on wave dynamics.

Furthermore, considering the role of the pulvinar in generating spontaneous FW waves may offer a potential explanation for the between-subjects variability we observed in experimental data at rest in the ratio between FW and BW waves. Unspecific attention or general arousal may modulate pulvinar engagement in this state and lead to a higher or lower probability of FW wave patterns at the scalp-level. Interestingly, in this case, our model would predict that the spatial distribution of these spontaneous FW waves should be more towards frontal regions, as compared to stimulus-evoked waves. With a suitable resting state paradigm, this could be tested in future studies.

It is important to notice that our investigation of the pulvinar pathway leaves out several notable properties of the thalamo-cortical connectivity. First, the pulvinar nodes in our model only passively relay activity, while physiological thalamic relay cells form a highly specialized circuit with the thalamic reticular nucleus that together exhibits complex non-linear response properties (Destexhe et al., 1998). Moreover, cortico-thalamic projections also comprise two distinct subtypes (drivers and modulators), and the interaction between these appears to play a role in the generation of alpha oscillations in the pulvinar itself (Abbas Farishta et al., 2020; Cortes et al., 2021; Grieve et al., 2000; Sherman, 2007). Lastly, our model does not include the backward-directed connection from one cortical area through the pulvinar to a lower area (see Shipp, 2003). This pathway is unlikely to assume a modulating role for BW waves equivalent to the one predicted for FW waves in our model, given that pulvino-cortical projections terminate largely in superficial layers (Kaas & Lyon, 2007; Ogren & Hendrickson, 1977). However, pulvinar input has been shown to play an important role in refining representations in V1 and gating its output (Purushothaman et al., 2012). This makes the pulvino-cortical feedback pathway a likely candidate for carrying prediction signals in the context of predictive coding (Cortes et al., 2024; O’Reilly et al., 2021). It will be important to consider this pathway in future implementations that expand the current dynamical framework to a feature space.

## Conclusion

In this work we presented a multiscale mean-field network model to explain and simulate the dynamics of alpha-band traveling waves measured with the EEG. We proposed two distinct inter-laminar pathways for the propagation of forward and backward alpha waves, and we modeled the role of the pulvinar in modulating the cortical pathways. Importantly, we ground the proposed architecture and the results of our simulations in the theoretical framework of predictive coding, in line with previous work. All in all, our model provides a first theoretical framework connecting these scalp-level waves to cortical circuits and could provide a base architecture for future research in several directions.

## Material & Methods

### Model Dynamics

Our model is based largely on the mean-field model introduced by Jaramillo et al. (2019). We used the same dynamics to model all cortical and thalamic nodes and their connectivity, except for the infragranular pacemaker (IG_IB_) described by average single-neuron dynamics. In the following, we first describe the regular mean-field dynamics and then extend it to the special case of the IG_IB_.

Each node *i* is described by its firing rate *r_i_* varying over time as a fixed function of the input current *I_i_* (FI-curve):

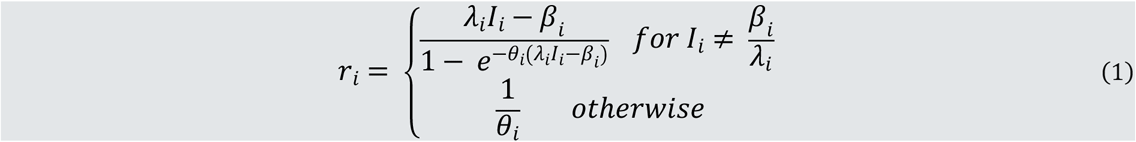

where r is in Hz, I is in nA, and λ, θ, and β are the FI-curve’s slope (excitability) and offset (neural threshold) parameters. The input current *I_i_* to the node is given by:

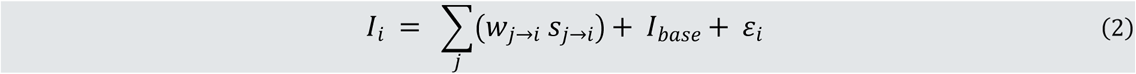

where the first part is the sum of the weighted synaptic input to node *i* (dynamics described below), *I_base_* is a constant base current, and ε_i_ is time-varying noise specific to each node. The noise term is an Ornstein-Uhlenbeck process of the form:

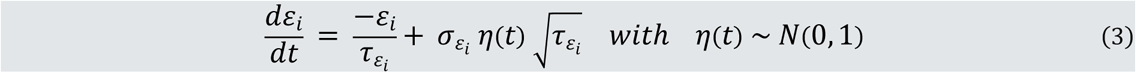

where τ_ε_ and σ_ε_ are parameters for the time constant and the strength of the noise, respectively.

A synapse *k* transmitting activity of node *i* to another node is defined through a constant weight *w_k_*, and a dynamic variable *s_k_* representing the gating of transmitter-mediated current flow, defined by the following dynamics:

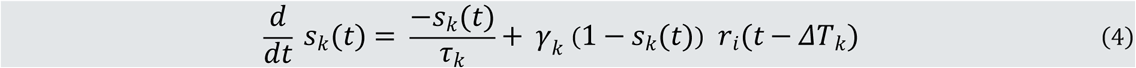

Where t is time, ΔT_k_ is the synaptic delay, *τ_k_* is the time constant of the synapse, *γ* ∈ [0, 1] is a saturation parameter and *r_i_* is the firing rate of node *i* as defined in (1).

### Infragranular pacemaker (IG_IB_)

To model the activity of intrinsically bursting populations in deep cortical layers, we substituted the mean-field response dynamics defined in (1) and (2) by a spiking-neuron model. For each node, we defined a homogeneous population of N = 350 Izhikevich spiking neurons (Izhikevich, 2003). Every neuron is described by two dynamic variables - its membrane potential *v* and a recovery variable *u,* which evolve according to the following equations:

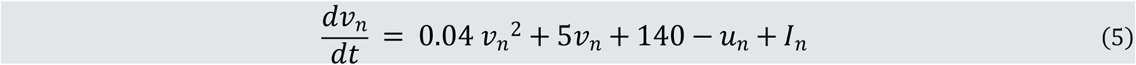

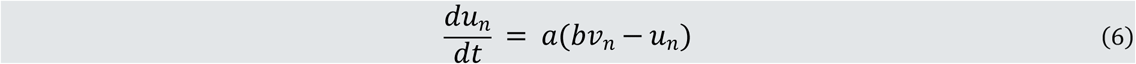

Here, *a* and *b* are parameters determining the dynamic behavior of the membrane potential (see Izhikevich, 2003). Once the potential passes a threshold, a spike is elicited and the values of *v* and *u* are reset:

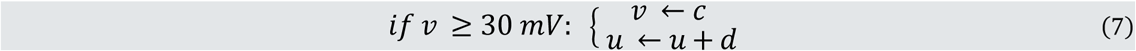

where c and d are the reset parameters. The values for a, b, c, and d were chosen such that the neurons generate short bursts of spiking activity at rest in regularly paced intervals of approx. 100ms (see list of parameter values below).

The activity of the spiking-neuron nodes was integrated into the mean-field model by applying fixed conversions at input and output stages (i.e., substituting equations (1) and (2) above). For a given spiking node *i_SN_,* its mean activity r_i[SN]_ is given by the instantaneous spike rate within the population:

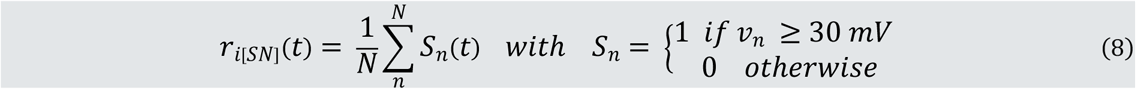

where N is the number of neurons in the population and *v_n_* is the membrane potential of neuron n as given by (5) – (7). As input, each neuron in the population receives the (scaled) input current to the node, plus a Gaussian noise term (independent for each neuron):

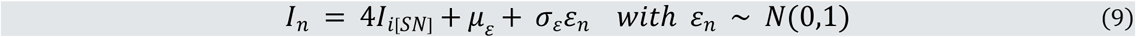

where *I_i[SN]_* is the summed input current to the node as defined in (3), and μ_ε_, σ_ε_ are the mean and standard deviation of the noise. Since the parameters *a-d* were identical between all neurons, the noise term in (9) is the only source of variability in the population. Note also that the neurons were not connected directly to each other. However, spiking nodes as a unit formed self-connections in the model (at the level of the mean-field input and output, i.e., equations (2) and (8)). This was designed to retain synchronization of the intrinsic bursting behavior of the population as a whole.

All equations of the model were evaluated using Euler’s method in time steps of 1 ms resolution, with the exception of the spiking neurons’ membrane potential ((5) and (6)) for which a 0.5 ms step was used.

### Cortical model architecture and connectivity

The general architecture and connectivity of the network are depicted in Figure 1. Our model describes a single-stream hierarchy of *N_Cx_* (visual) cortical areas. Unless noted otherwise, all simulations use a version of the model with *N_Cx_* = 3. The general response behavior and dynamic reversal of traveling wave direction remained the same when we tested versions with up to 7 areas.

Each cortical area comprises six nodes across three laminar compartments: L4_X_ and L4_IN_ in Layer 4, SG_X,_ SG_E,_ and SG_IN_ in supragranular Layers 2/3, and IG_IB_ in infragranular Layers 5/6, where the *IN* subscript denotes inhibitory nodes. We focus our analysis mainly on SG_X_ and IG_IB_ activity as the central nodes in the FW- and BW-pathways (see main text).

Our choice of model architecture was motivated in equal parts by the hierarchical predictive-coding model developed by Alamia & VanRullen (2019) and physiological evidence of functional connectivity between laminar compartments. The predictive coding framework (Rao & Ballard, 1999) postulates that feedforward projections mainly carry the difference (prediction error) between the information of the sending area and that sent as feedback from the receiving area (prediction). In our model, the prediction error is explicitly encoded by the SG_E_ unit in each area. We describe how our architecture relates to the original model (Alamia & VanRullen, 2019) in the supplementary material (S1).

As a reference for laminar connectivity (as well as existing neural implementations of predictive coding), we refer to previous work by Shipp et al. (Shipp, 2016; Shipp et al., 2013) and Bastos et al. (2012). Our architecture follows the dominant flow of information in the canonical microcircuit from Layer 4 to superficial and from there to deeper layers. We also assume that inter-areal feedforward connections project mainly to Layer 4 while feedback projections terminate in supra- and infragranular layers (Markov et al., 2014).

The input current in our model is applied to a separate input layer (IL_St_) comprising one pair of recurrently connected excitatory and inhibitory nodes (identical to Layer 4 of each cortical area). This stage was included for additional input normalization prior to Cx_1_, and may be seen as equivalent to LGN.

An identical pair of nodes was included at the other end of the network (IL_Pr_). The excitatory node in this pair was connected to the IG_IB_ node in the last cortical area (Cx_N_) and received a continuous white noise input (varying between 0 and 0.3 nA). This was not intended to model a prior signal (as in Alamia & VanRullen, 2019), but it merely acts as a cortical top-down signal, ensuring stable activation of the BW-pathway.

Unless stated otherwise, cortical synaptic delays were fixed at 0 ms for local (intra-areal) and 12 ms for remote (inter-areal) connections. It should be noted here that the ‘net’ (e.g., as peak-to-peak) delay for activity sent from one node to another depends on the synaptic delay, synaptic time constant and neural integration times of the receiving and any intermediate nodes.

### Pulvinar pathway

The thalamocortical version of our model includes, in addition to the cortical modules, a single pulvinar node between any two consecutive cortical areas, modeling the functional feedforward connectivity of thalamic relay neurons. The connection delay between cortex and Pulvinar was fixed at half the cortical inter-areal delay (i.e., 6 ms). That is, the summed delay of the connections Cx-Pul-Cx was the same as that of direct Cx-Cx connections. However, we confirmed that the main pattern of results for the pulvinar simulations were the same with double delays.

### EEG simulation

We used a simple forward model to obtain a scalp-level projection of our model’s mean-field output. The primary goal of this was to compare wave dynamics at the laminar level in our model to existing EEG data, using a phase-based wave fitting procedure (see below).

The activity for each cortical node was first low-pass filtered using a 3^rd^ order Butterworth filter (cutoff 20 Hz) and then down-sampled to 100 Hz from the 1kHz sampling rate of the simulations. For each simulated trial, a set of 5 independent noise sources (power spectrum slope 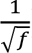) were generated and projected together with the laminar signals. The signal-to-noise ratio (with respect to the root mean square of both signals) for each noise source was drawn randomly from a uniform distribution between 0.4 and 1.6.

The forward model projections were performed using functionalities provided by the fieldtrip toolbox (*ft_dipolesimulation*; Oostenveld et al., 2011). Source dipoles were oriented radially and positioned according to their area, using the following coordinates in MNI space (based on the AAL atlas, Tzourio-Mazoyer et al., 2002):

**Table 1.** Coordinates of dipole positions used for the EEG forward model. All values are MNI coordinates based on the AAL atlas (Tzourio-Mazoyer et al., 2002).

| *Model Area* | *Label* | *Coordinates* |
| --- | --- | --- |
| Cx<sub>1</sub> | Occipital | Left: [-8 -76 10]<br>Right: [8 -76 10] |
| Cx<sub>2</sub> | Parietal | Left: [-24 -61 58]<br>Right: [24 -61 58] |
| Cx<sub>3</sub> | Frontal | Left: [-5 48 30]<br>Right: [5 48 30] |

For the single-stream version of the model (used everywhere except in the simulations of hemispheric lateralization), the cortical sources were positioned bilaterally in both hemispheres, scaled at half amplitude each. The noise sources were positioned randomly for each trial on a 5 x 5 mm grid covering all positions located inside the skull. Neither the pulvinar nor the stimulus input layer (IL_St_) were included in the forward projection.

### EEG data collection and analysis

The EEG data used in the comparisons with our model output have been reported in more detail previously (Pang (庞兆阳) et al., 2020). In brief, observers (*N = 13*) were presented with a single, iso-luminant disk (7° diameter, centered at 7.5° above fixation) for a duration of 5 sec on each trial, followed by 5 sec of blank screen. The disk’s luminance either followed a white-noise random sequence (“Dynamic” condition) or remained uniform (“Static”). We include only the “Static” condition in our comparison. Throughout all trials, subjects performed an adaptive visual detection task requiring them to attend the stimulus covertly.

All pre-processing steps for the present analyses were identical to the previous study. Signals were recorded on a 64-channel BioSemi system at 1024 Hz and down-sampled offline to 160 Hz temporal resolution. Re-referencing was performed by subtracting the average activity and band-stop (50 Hz) and high-pass (> 1Hz) filters applied. Artifact detection and rejection were performed manually on the epoched data.

### Wave fitting procedure

To quantify the direction of wave propagation in both simulated and real EEG signals, we adapted the procedure introduced by Zhang et al. (2018) for use with scalp-level data. This method relies solely on the analytic phase of the signal and is therefore well suited to analyze our simulated data (in contrast with the 2D-FFT method used in previous studies (Alamia et al., 2023; Alamia & VanRullen, 2019) that also factors in amplitude topography which was not the target of our simulations).

For a given frequency band of interest (7-13 Hz unless otherwise stated), the continuous phase of the signal was obtained using a bandpass FIR filter and the analytic Hilbert transform. The resulting phase values were referenced to the mean phase within the electrode ROI at each timepoint. The following iterative fit procedure was evaluated using the time-averaged relative phase within a sliding window 100 ms. For all analyses, we used a 2D projection of the electrodes’ positions on the scalp (*10-20* topography layout provided by the fieldtrip toolbox; Oostenveld et al., 2011).

For a set of electrode positions x, y, the phase values 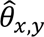 predicted by a perfect planar traveling wave are given by:

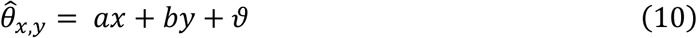

where a, b are free parameters and ϑ is the reference-(mean) phase. The propagation direction of the wave is given by α = *atan*2(*b, a*) and its spatial frequency by 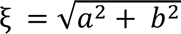.

For each time-point, the model was evaluated iteratively within a given set of values for a and b. This parameter space was defined by 60 steps over the possible space of wave directions ξ (360°) x 30 steps of increasing spatial frequency up to 1 cycle over the maximum distance covered by the electrode space. The best fit was determined by finding the maximum vector length of the summed residuals in circular space:

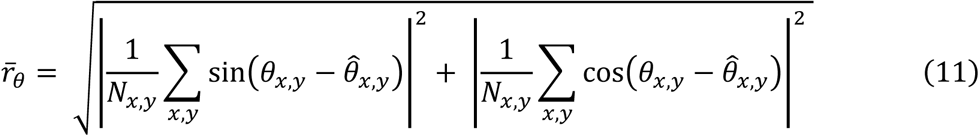

Further following Zhang et al. (2018), the circular correlation ρ between predicted (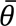) and observed (θ) phases was used as a statistical measure for the absolute goodness of fit:

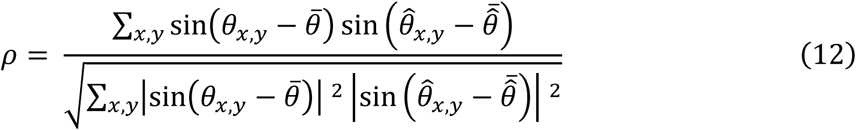

with 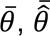 denoting the mean within the ROI of observed and predicted phases, respectively. To determine the significance of a given fit, we compared it to a chance-level distribution of ρ values (using a one-sided threshold of 95%). This distribution was obtained once for each dataset by repeating the fitting procedure 10 times with randomly permuted electrode positions, at reduced temporal resolution.

Finally, each time-point was classified into one of three categories: as “Forward” (“Backward”) state, if the fit was statistically significant and the propagation direction was within 0.5 rad of the FW/BW axis (see below), and as a “Null” state otherwise. For non-lateralized stimulation, activation can be assumed to be symmetrical between hemispheres on average therefore the reference axis is defined as the

Fronto-Occipital axis. Spatial propagation in response to lateralized stimulation can deviate from this axis (Alamia et al., 2023; Lozano-Soldevilla & VanRullen, 2019). For lateralized simulations, we therefore defined the reference as the median propagation direction after combining BW and FW directions into a single distribution through collapsing on the medio-lateral axis.

### List of parameter values

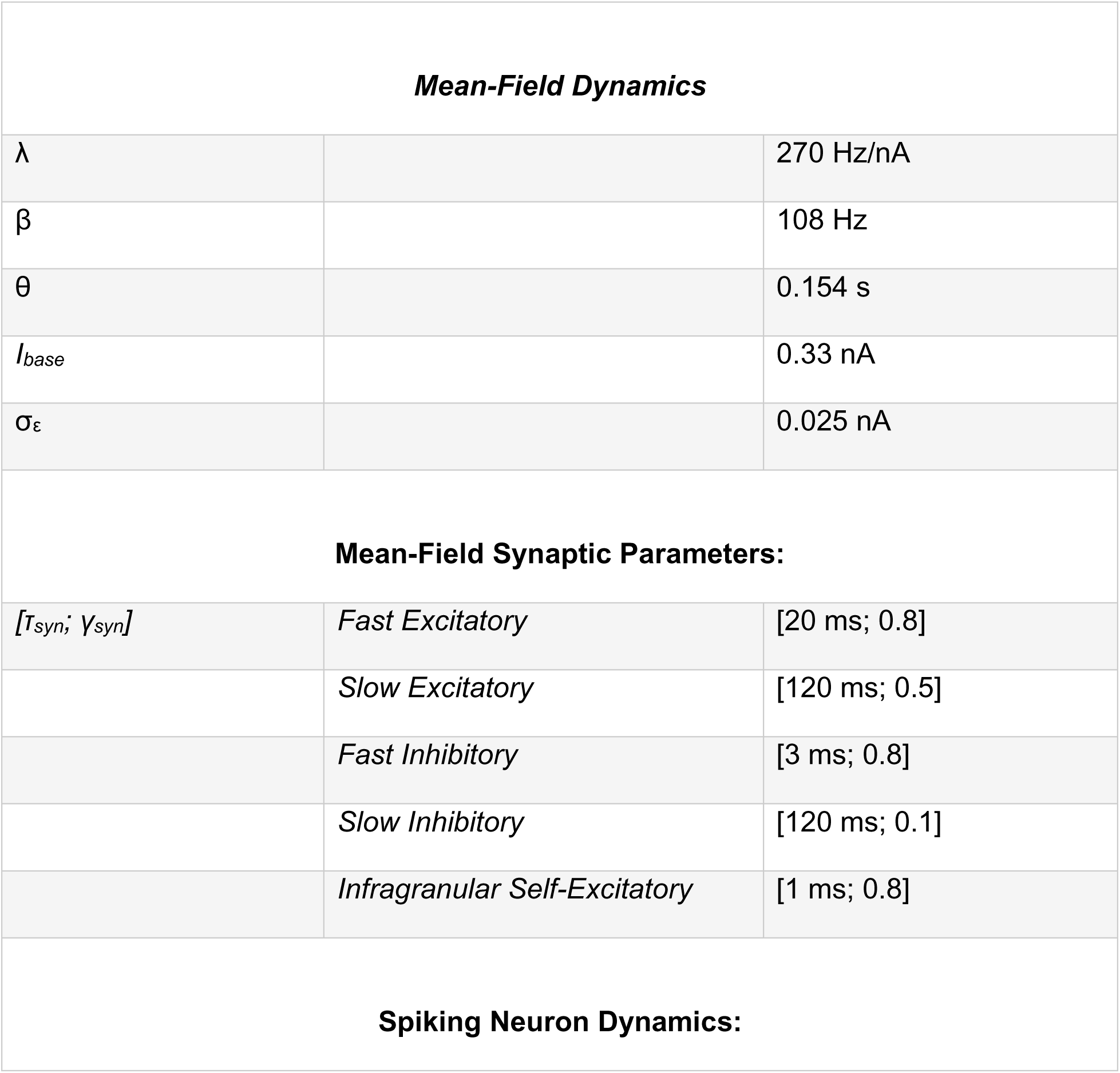
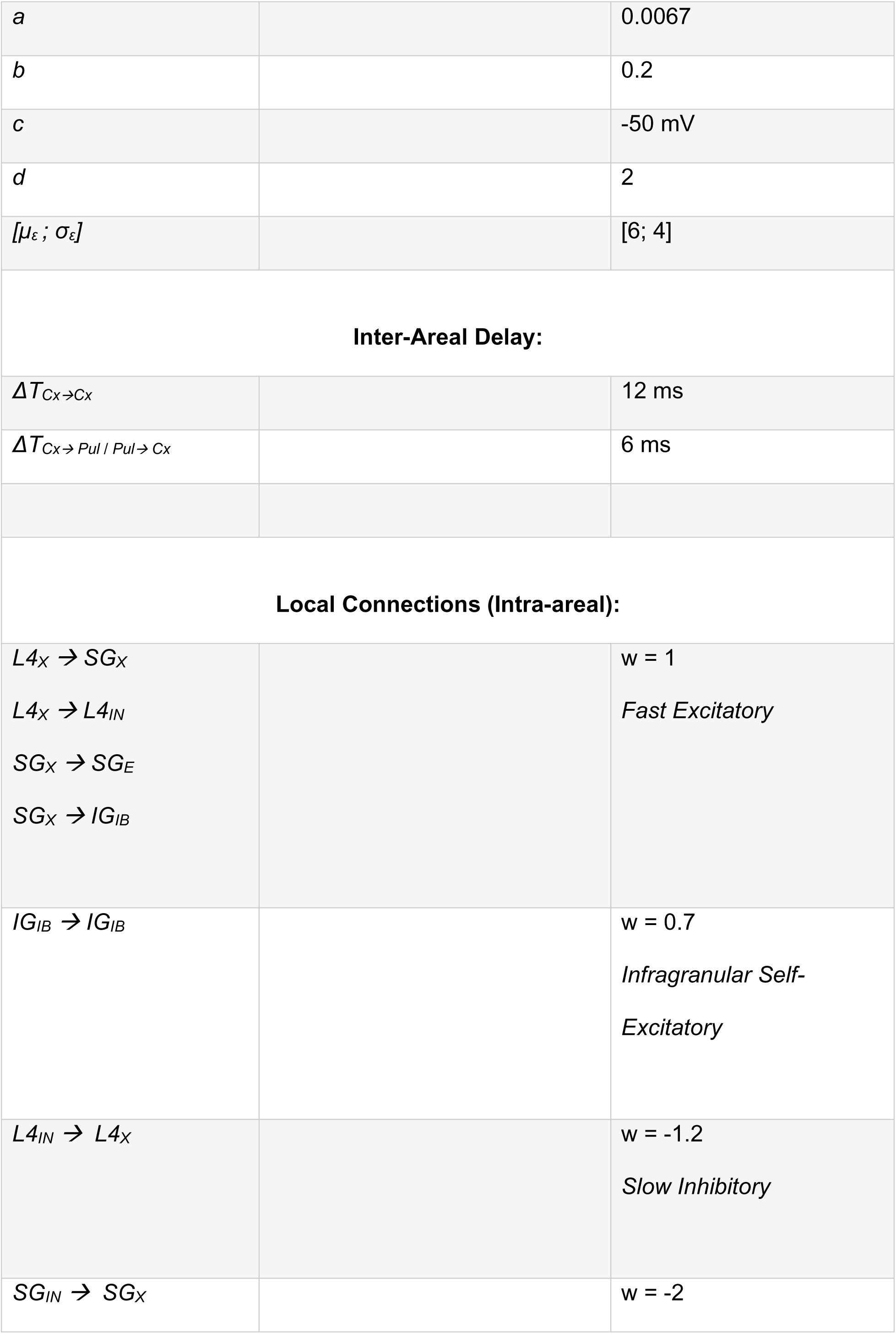
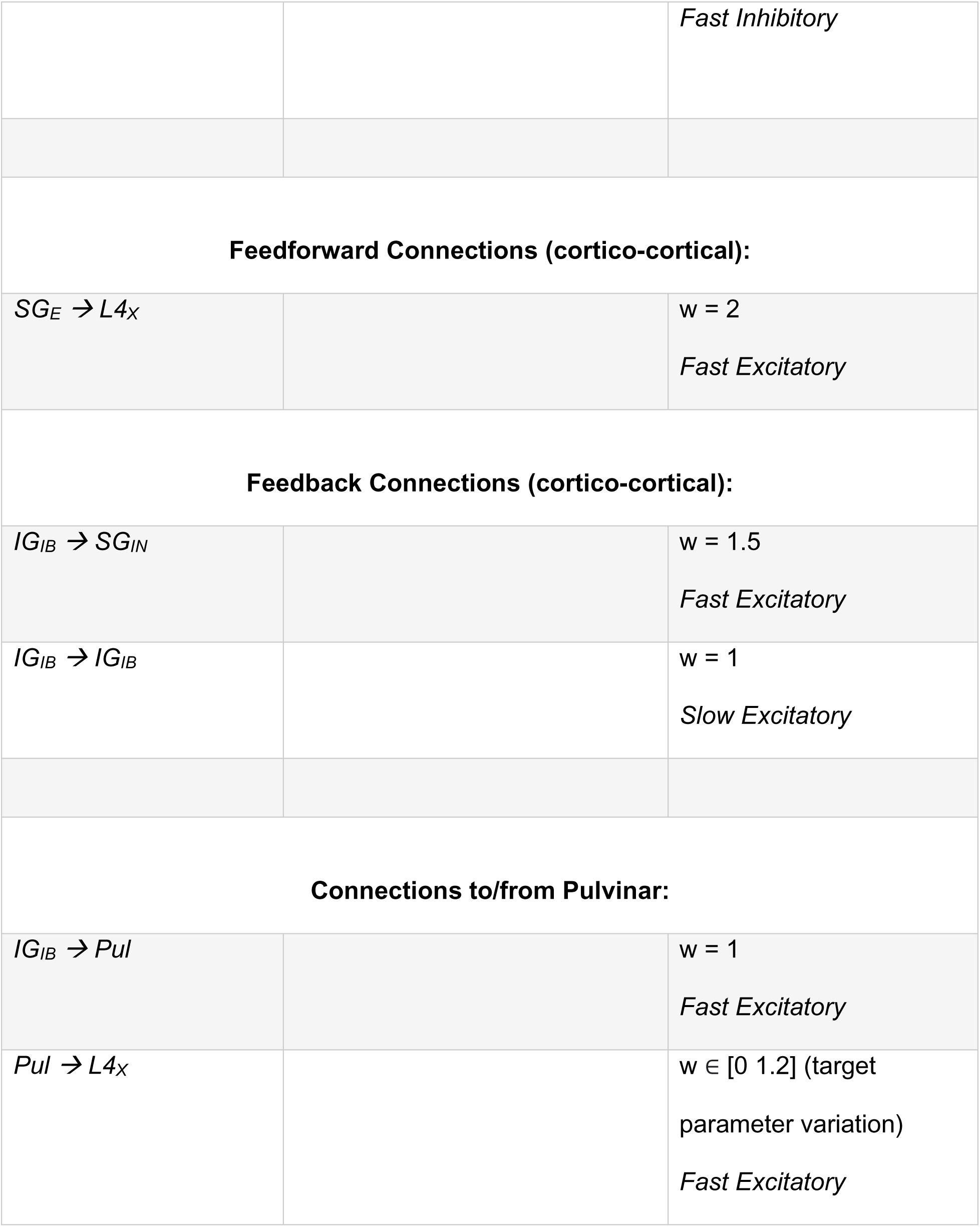

## Acknowledgements

This project was funded by the European Union under the European Union’s Horizon 2020 research and innovation program (grant agreements No. 101075930 to Andrea Alamia). The copyright holder for this those of the author(s) only and do not necessarily reflect those of the European Union or the European Research Council (ERC). Neither the European Union nor the granting authority can be held responsible for them. The authors would like to thank Xiaoqi Xu and Martin Vinck for helpful comments on the model, as well as Leslie Marie-Louise for administrative support.

## Notes

### Competing Interest Statement

The authors have declared no competing interest.

