## Supplemental Material S1 for "A hierarchical multiscale model of forward and backward alpha-band traveling waves in the visual system"

*Derivation of the FW-pathway architecture in our model from the hierarchical model in Alamia & VanRullen (2019)*

The circuit generating FW waves in our model (FW-pathway) is based on the predictive coding (PC)-model developed by Alamia & VanRullen (2019). Here, we provide in more detail our reasoning in choosing the architecture for this part of our model.

The PC-model is explained in more detail in the original publication. Briefly summarized, it comprises a hierarchy of layers, in which each layer represents a continuous prediction about the activity of the previous layers. Feed-forward signals contain the prediction error, i.e., the difference between prediction and the received input. As a dynamical system, for layers *L = 1 … N*, this is defined through the following equations:

$$x_L(t) = y_{L-1}(t) - y_L(t - \Delta T) \tag{1}$$

$$\frac{dy_L}{dt} = \frac{1}{\tau} x_L(t - \Delta T) + \frac{1}{\tau_D}\left(y_{L+1}(t - \Delta T) - y_L(t)\right) \tag{2}$$

where $y_L$ is the prediction at layer L, $x_L$ is the prediction error forming the input into layer L, t is time, ΔT is the temporal delay between layers, and $\tau$ and $\tau_D$ are time constants for the integration of incoming signals and the decay of local signals, respectively. Plausible values for the temporal variables are $\tau$ = 20ms, $\tau_D$ = 200 ms and ΔT = 12 ms (see Alamia & VanRullen, 2019, for an exploration of the parameter space).

To derive a physiological implementation of this model, we first put the above into a neural framework (Figure 1). The continuous prediction signal y can be seen as the local stimulus representation in each area. Eqs. (1) and (2) include two key computations that also occur locally. First, the prediction error x is computed by subtracting the top-down prediction from the local one (eq. (1)). This is achieved by a second node that represents x, receiving at equal weights excitatory input from the local- (at zero delay) and inhibitory input from the top-down prediction (at delay ΔT). Note that the indexing of x is somewhat arbitrary and may either be based on the layer at which is it computed or on the one it is fed forward to; Figure 1 uses the latter, same as eqs. (1) and (2). Secondly, the local prediction y is slowly (large time constant $\tau_D$) updated by comparing itself to the top-down prediction (second part of eq. (2)). This is equivalent to simultaneous decay of the prediction and an excitatory input from the top-down prediction at equal weights. Lastly, the feed-forward prediction error is weighted by the time constant τ.

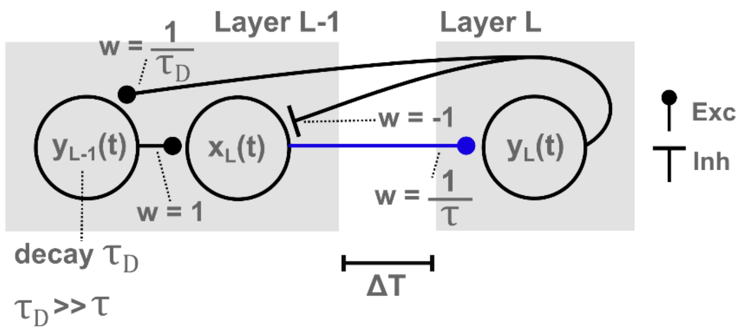


*Figure 1. Neural node-graph representation of the original PC-model. Variable and parameter names correspond to eqs. (1) and (2).*

From this basic architecture, we derive the architecture of our physiological model as follows. First, we expand each area by an input layer L4 which relays the feed-forward connection (blue) (Figure 2). Representation and error nodes are re-labeled as $SG_X$ and $SG_E$, respectively, to reflect their location in the supragranular layers. The excitatory input from the top-down prediction (dashed line in Figure 2) in the original PC-model allows for a backward propagation of top-down signals through the network. Since we split up FW and BW propagation into separate pathways, this connection is eliminated here (it is partly represented in the full model by the infragranular connectivity of the BW-pathway).

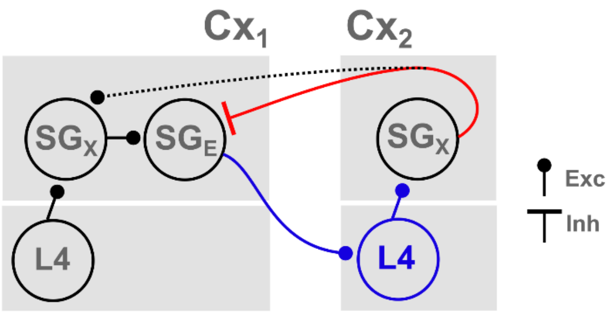


*Figure 2. Expansion of the PC-model graph into a laminar structure. The feed-forward connection (blue) is now relayed through layer 4. The excitatory feedback carrying the top-down prediction (dashed) is eliminated in the FW-pathway.*

Finally, for physiological plausibility we add a local inhibitory node to mediate the inhibitory feedback (red), and extend it to a more plausible pathway through the infragranular layers (Figure 3). This architecture corresponds to the FW-pathway in our model. It satisfies the main physiological boundary conditions, and matches with the functional microcircuits proposed by others as laminar implementations of predictive coding (see, e.g., Shipp, 2016; Shipp et al., 2013). Note that it relates to the original PC-model mainly in its inter-areal connectivity (including the inter-areal delay), whereas neural dynamics of the individual nodes are based on a different model (see main paper).

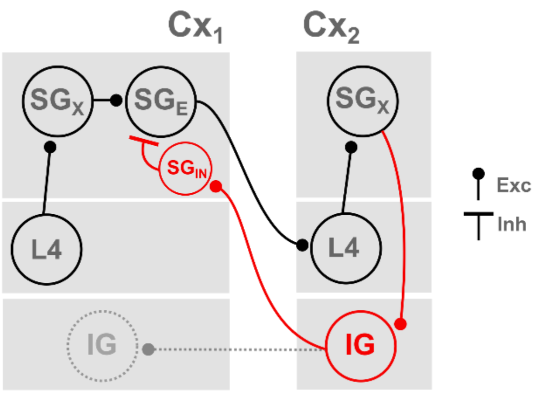


*Figure 3. Final FW-pathway from the physiological model. The inhibitory feedback for the computation of the prediction error (red) is now relayed through infragranular layers. In the*

*full model (see main paper), the IG nodes are intrinsically bursting with backward coupling between areas (BW-pathway, dashed).*
